# 3D super-resolution imaging of PSD95 reveals an abundance of diffuse protein supercomplexes in the mouse brain

**DOI:** 10.1101/2024.10.14.617231

**Authors:** Sam Daly, Edita Bulovaite, Anoushka Handa, Katie Morris, Leila Muresan, Candace Adams, Takeshi Kaizuka, Alexandre Kitching, Alexander Spark, Gregory Chant, Kevin O’Holleran, Seth G. N. Grant, Mathew H. Horrocks, Steven F. Lee

**Author notes:** S.D. and E.B. contributed equally to this work.

## Abstract

PSD95 is an abundant scaffolding protein that assembles multiprotein complexes controlling synaptic physiology and behavior. Confocal microscopy has previously shown that PSD95 is enriched in the postsynaptic terminals of excitatory synapses and 2D super-resolution microscopy further revealed that it forms nanoclusters. In this study, we utilized 3D super-resolution microscopy to examine the nanoarchitecture of PSD95 in the mouse brain, characterizing over 8 million molecules. While we were able to identify structural subtypes previously reported, imaging in 3D allowed us to classify these with higher accuracy. Furthermore, 3D super-resolution microscopy enabled the quantification of protein levels, revealing an abundance of PSD95 molecules existed outside of synapses as a diffuse population of supercomplexes, containing multiple copies of PSD95. Further analysis of the supercomplexes containing two units identified two populations: one that had PSD95 molecules separated by 39 ± 2 nm, and a second with a separation of 94 ± 27 nm. These results suggest that PSD95 supercomplexes containing multiple protein copies assemble outside the synapse and then integrate into the synapse to form a supramolecular nanocluster architecture.

## Introduction

Synapses are specialized junctions between nerve cells that contain thousands of proteins and underpin our understanding of cognition and learning^1–5^. Most synaptic proteins are organized into multiprotein complexes that are essential for key functions such as synaptic transmission and plasticity^6^. Biochemical studies confirm that scaffolding proteins assemble a variety of neurotransmitter receptors, ion channels, signaling proteins and transmembrane proteins into families of complexes in the postsynaptic terminal of excitatory synapses^6,7^. Classical electron microscopy studies have shown an electron-dense structure beneath the postsynaptic membrane of excitatory synapses, known as the postsynaptic density (PSD)^8–10^. Recent studies employing cryogenic correlated light-electron microscopy (cryoCLEM) demonstrate that, rather than an increase in postsynaptic protein density, these molecules exhibit a specialized organizational structure in the postsynaptic terminal^11^.

The prototypical scaffolding protein of excitatory synapses is postsynaptic density protein 95 (PSD95), which plays a crucial role in synaptic transmission, synaptic plasticity and learning^12^. PSD95 is highly abundant within synaptic protein extracts and found within 1–3 MDa complexes^7,13,14^. Confocal microscopy of labeled endogenous PSD95 has confirmed its presence in the postsynaptic dendritic spine^15–17^ and super-resolution (SR) microscopy has explored the nanoscale architecture of PSD95, and other proteins in synaptic clusters, in cultured cells^18,19^ and brain tissue^20–22^. These studies revealed that approximately 100 PSD95 molecules were organized into nanoclusters (NCs)^14,20^ around 130 nm in size^20^ and located in the PSD, which varies in size from ∼100–500 nm in diameter^20,23^. Receptors, including AMPARs have also been shown to form similar nanoclusters^19^ and their functional relevance has been demonstrated *via* their alignment to clusters at the presynapse, by so-called “nanocolumns”^24^. The prevailing model suggests that PSD95 molecules participate in an assembly process where supercomplexes containing two copies of PSD95 are formed, which we have previously characterized^25^. These supercomplexes then assemble into larger multiprotein complexes, which ultimately combine into NCs to form the PSD^26^. However, it is still unclear whether all these assembly stages occur within the synapse itself, which limits our understanding of how synaptic structure and function are regulated both spatially and temporally.

To understand the organization and assembly of PSD95 into supramolecular structures, we have developed various genetic labeling strategies for SR microscopy^20,21,27^. In particular, single-molecule localization microscopy (SMLM) has shown that synapses contain multiples of NCs (1 NC/PSD, 2 NC/PSD and 3+ NC/PSD) that are differentially distributed into synapses in regions of the hippocampal formation—a brain region important for learning and memory^20^. However, this SMLM approach was limited to visualizing PSD95 in two dimensions and in regions close to the cutting plane of the fixed brain slices, which may have been subject to dissection artifacts^28^. Furthermore, imaging biological samples in 3D is crucial to avoid clustering artifacts that can distort quantitative imaging results and lead to inaccurate interpretations of spatial organization.

Various methodologies are available for studying protein distribution at the nanoscale, each offering different depth-of-field (DoF) and resolution specifications that depend on the sample being studied. These include multifocal plane approaches, such as biplane microscopy (<1 µm DoF)^29^, and point spread function (PSF) engineering techniques. PSF engineering methods include astigmatism (<1 µm DoF)^30^, the double-helix PSF (DHPSF, 2–4 µm DoF)^31^, the tetrapod PSF (4–8 µm DoF)^32,33^, and single-molecule light field microscopy (∼8 µm DoF)^34,35^. The DHPSF is well-suited to the investigation of sectioned tissue by affording greater resolution isotropy (xy: ∼10 nm and z: ∼20 nm)^36^ compared to astigmatism^37^ and a large ∼4 µm DoF that eliminates the need for axial scanning^38^.

We present the first application of DHPSF SR microscopy in tissue, enabling the most comprehensive study of the 3D organization of individual PSD95 molecules. We evaluate 8,407,417 single PSD95 proteins, and 24,931 NCs in 7 brain regions from 3 mouse brains. Not only do our results support previous findings from 2D SR imaging regarding the spatial architecture of PSD95 within synapses, but the capability to image tissue within the bulk section led to the discovery of an undescribed subpopulation. We reveal that the majority of PSD95 exists as a ‘diffuse’ cloud of complexes containing multiple copies of the protein throughout the tissue volume, rather than being associated with the synapse.

## Results

### Volumetric SR imaging of PSD95 NCs using DHPSF microscopy

A functional advantage of SMLM is its ability to resolve synaptic diversity across increasingly complex spatial scales, beginning with individual proteins, their assembly into NCs, the formation of building blocks within PSDs, and ultimately their differential spatial arrangement across the hippocampus (see **Figure 1**). Building on our previous genetic labeling strategy^39^, we imaged intact mouse brain tissue samples of mEos2-labelled PSD95 using DHPSF SR microscopy (see **Figure S1 & SI Movie 1**). The DHPSF enabled the determination of the true Euclidean coordinates (x,y,z) of each PSD95 molecule with nanometer accuracy. These volumetric pointillist images correspond to the sub-diffraction distribution and organization of PSD95 (see **SI Movie 2**). Like our previous study in 2D^20^ clustering algorithms allowed us to segment the localization data to distinguish thousands of PSDs composed of individual NCs across a large field-of-view in 3D (70 × 70 × 4 µm, see **Figure 2a & Methods**). Furthermore, our analysis of mEos2 single-molecule photophysics mitigated the risk of molecular overcounting and thus validated our quantitative approach to measuring protein spatial distributions (see **Methods** and **Figure S2** for further details).

**Figure 1.**
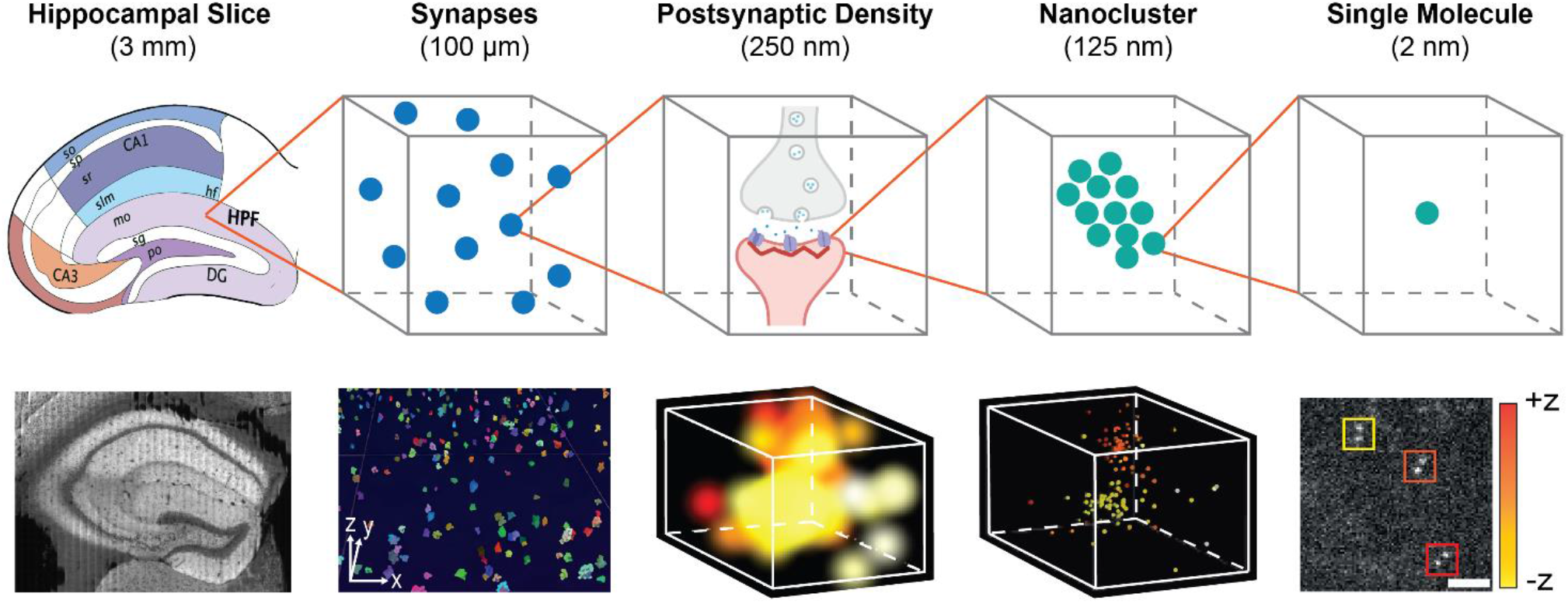
PSD95 Assembly at different spatial scales. (Top) Schematic representations and relative sizes of the coronal hippocampus, a synapse, postsynaptic density (PSD), a nanocluster (NC) of PSD95, and a single mEos2-labelled PSD95 molecule. (Bottom) Raw data corresponding to the schematic shown above. Hippocampal slice: confocal image of a coronal hippocampus mouse brain slice. Synapses: CA1SR hippocampal region (30 × 30 × 4 µm) with ∼200 NCs; PSD: diffraction-limited image of PSD95 NCs; Nanocluster: super-resolved PSD95 NCs; Single molecule: DHPSF SR microscopy data of a single PSD95 molecule fused to the fluorescent protein mEos2. Color indicates axial position and scale bar represents 5 µm.

**Figure 2.**
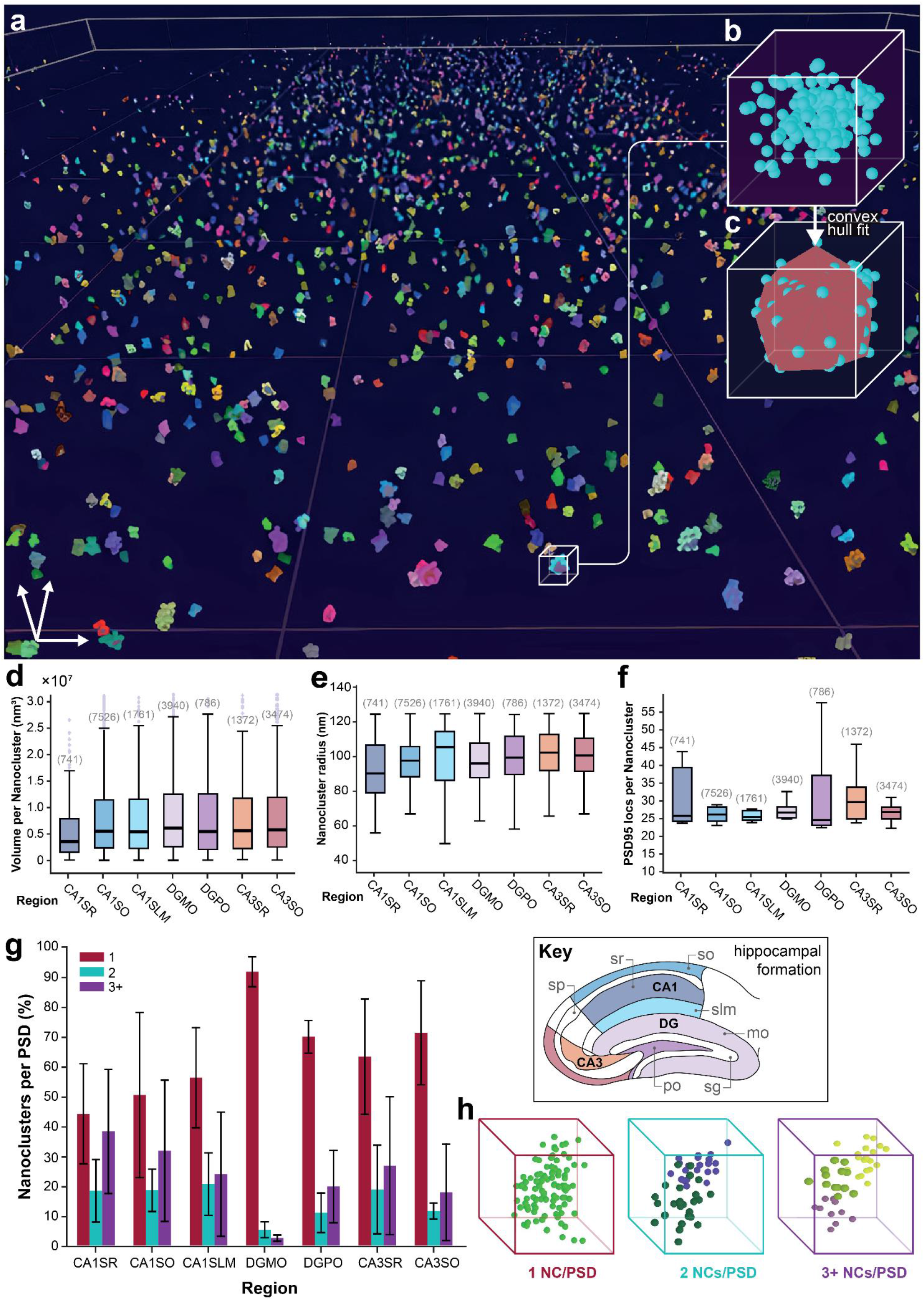
3D DHPSF imaging of PSD95 NCs. **a** Representative reconstruction of the CA1SR region (70 × 70 × 4 µm) with NCs represented as demarcated colors. **b** Expanded view of a NC with ∼30 localizations of PSD95. **c** Convex mesh fitting to the NC from b) used to calculate the volume of NCs. **d** Volume per NC by region calculated using the triple product convex mesh-fitting technique (sample size of NCs given above each box plot). A depiction of brain regions imaged in this work color-coded for comparison with regional box plots (e–g) is presented below. **e** Radius per NC by region assuming them to be spherical. This data agreed with a complementary analysis of the radius by PCF (see **Figure S4**). **f** Mean number of PSD95 localizations per NC by region. **g** Number of NC/PSD as a percentage by region determined by Pair Correlation Fitting (PCF, mean ± 1 std). **h** Examples of 1 NC/PSD, 2 NCs/PSD and 3+ NCs/PSD. Visualization was carried out using vLUME desktop. Box plots represent the median and interquartile range, where whiskers extend to the extreme data points which are not considered to be outliers.

We first segmented the 3D localization data into NCs within PSDs using DBSCAN (cluster radius of 125 nm and a minimum size of 10 localizations, see **Methods**), thereby distinguishing between NCs within the same PSD and those in adjacent PSDs based on their spatial arrangement and distance. The addition of the z dimension better represented the true organization of NCs, and allows us to evaluate not only their radius but also their volume (see **Figure 2b & c**).

The 3D data can be contrasted with the 2D results in three distinct ways. First, we observe a decrease in the number of PSD95 localizations per cluster from ∼90 (in 2D)^20^ to ∼25 (in 3D). This can be attributed to a reduction in photon efficiency in the 3D method (∼30% loss^40^), which arises from to the incorporation of phase-modifying optics in the emission path. Second, we observe an increase in NC radii, which is justified by improved segmentation in 3D by avoiding reconstruction artifacts from the projection of a 3D volume onto a 2D plane. Finally, we observe an increase in the relative complexity of the PSD subtypes, which can also be attributed to improved 3D segmentation. For example, when observed in 2D, two NCs that are laterally aligned but axially displaced could be misclassified as a single NC per PSD, whereas 3D SR imaging allows for accurate distinction between them.

Analysis of NC populations across the hippocampus (see **Figure S3**) demonstrated that NC radii, volumes, and the number of localizations per NC were generally uniform across all regions examined, with values of approximately 100 nm, 0.5 nm3, and 25 localizations, respectively (see **Figures 2d–f & S4a**). Contrastingly, the PSD subtypes of 1, 2 and 3+ NC/PSD varied across the regions. Specifically, the proportion of 2 and 3+ NCs/PSD was typically higher in 3D, except for the DGMO region (see **Figures 2g & h**). For example, in the CA1 region, we see the ratio of 1:2:3+ NCs/PSD as 52:18:30, which contrasts with the ratio of 75:19:6 that was observed previously in 2D^20^. Taken together, these data confirm the presence of NCs *in vivo*, and their nature as highly conserved building blocks underpinning the structure of PSDs.

### Identification of a ‘diffuse’ PSD95 population

Segmentation-by-clustering allowed us to determine ‘hot-spots’ where localizations were at high density enabling us to distinguish PSDs (see **Figure 3a & Methods**). This approach typically removed randomly distributed localizations, such as noise arising from localization fitting errors, fluorescent impurities on the coverslip, and cutting artifacts to which 2D-SMLM is more sensitive^41,42^. However, in 3D these interfacial phenomena are not present as we are imaging throughout the entire tissue section and false positive localizations are much less common owing to the complex PSF shape (see **Figure S5**). The DHPSF therefore allowed us to probe the spatial distribution of all detected PSD95 molecules, not just those segmented in NCs. Upon reevaluating our 3D data, we unexpectedly discovered that most PSD95 are not confined to the PSDs but dispersed as a ‘diffuse cloud’ throughout the brain (see **Figure 3b & SI Movie 3**). A simple quantification of the number of the clustered and non-clustered localizations revealed that over 90% of protein is in fact not associated with NCs or PSDs, and this effect is broadly uniform over all regions examined (see **Figure 3c**).

**Figure 3.**
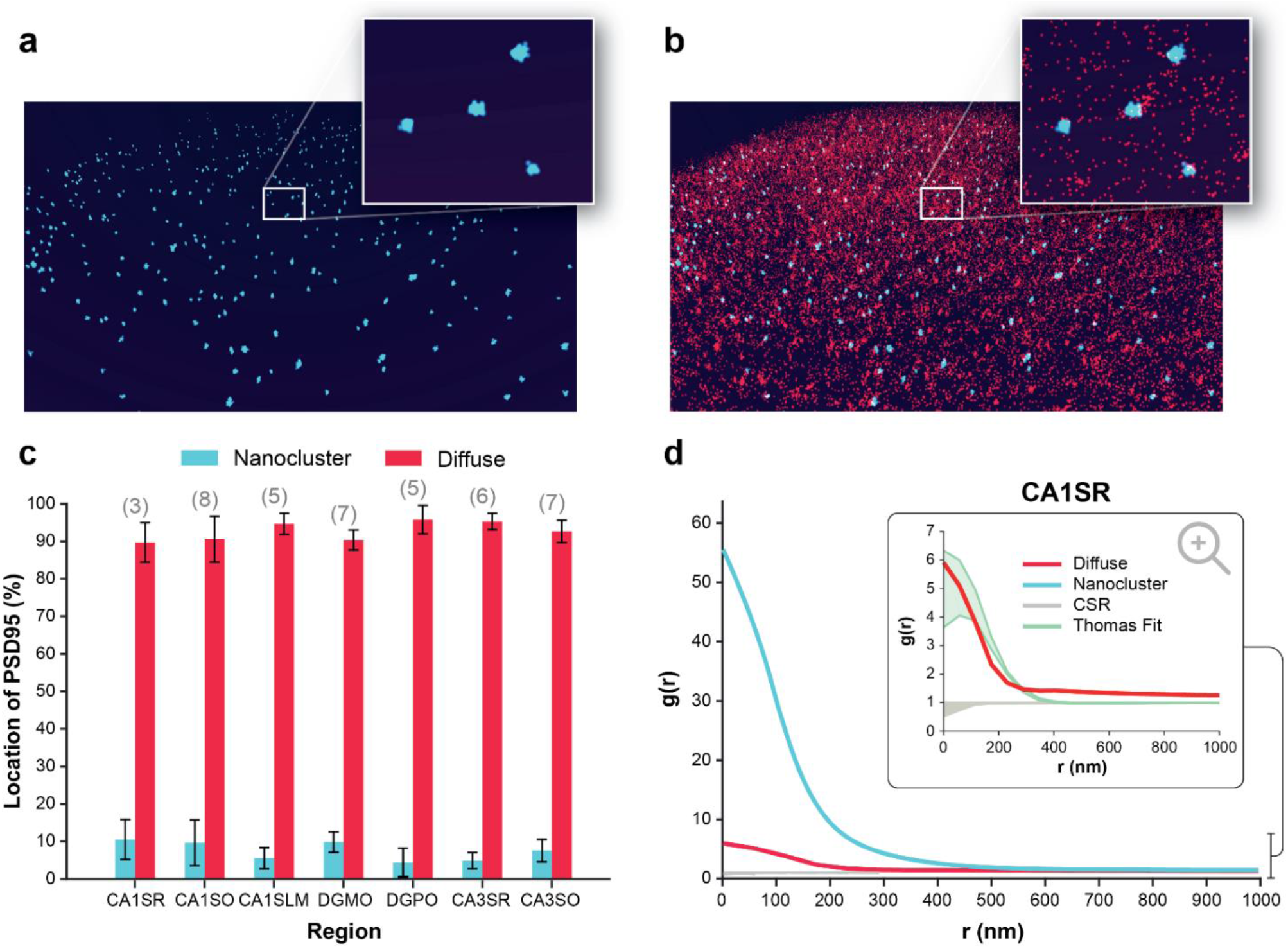
An abundance of PSD95 is found outside of the PSD. **a** Volumetric distribution of PSD95 nanoclusters (blue) in the CA1SR region. **b** Volumetric distribution of NCs and diffuse localizations (red). **c** Distribution of PSD95 localizations associated with the NC and diffuse populations. Error bars represent the standard deviation from the mean (bar) across the replicate tissue sections (shown above each region). **d** PCF analysis of the diffuse (red) and NC (blue) populations in the CA1SR region. This shows the difference in the degree of clustering of the two populations. The two envelopes correspond to the complete spatial randomness (CSR, gray) and Thomas fitting (green) simulations. The red line shows that the diffuse population does not adhere to a complete spatial randomness fit. The magnified region highlights the minor observed clustering present in the diffuse population. PCF analysis results from other regions are presented in **Figure S8**.

To confirm that this observation was not an artifact of the mEos2-labelled PSD95, we used genetically modified mice expressing eGFP^15^ or the HaloTag^16^ fused to PSD95. Spinning disk confocal microscopy was used to image the eGFP- or Halo-tagged PSD95 in mouse brain sections (see **Figure S6**). While both methods revealed an abundance of PSD95 outside of the PSDs, the percentage detected using spinning disk confocal microscopy was lower than that observed with mEos2.

To explore this finding in greater detail, we conducted a more sophisticated analysis approach. The localization data was once again segmented into clustered and diffuse datasets, this time using a custom k-Nearest Neighborhood (NN) algorithm (see **Methods**). Subsequently, pair correlation function (PCF) analysis was performed on each dataset (see **Figure 3d**), which represents the probability of encountering a second localization at a given distance relative to a reference point (normalized for the intensity/density of the overall localizations)^43^. For the clustered data, there is a high degree of correlation extending out to distances of ∼400 nm and the cluster radii corresponded well with our previous analysis approach (see **Figure S4**). On the other hand, PCF analysis of the ‘diffuse’ PSD95 population revealed greater disorder, but spatial correlation was still observed at smaller radii (up to ∼ 200 nm). For example, the g(r) values for NCs are ∼7-fold larger than the ‘diffuse’ population in the CA1SR region. To further validate these findings, localization data were simulated using a complete spatial randomness model, in addition to single and double Thomas fitting to generate parameters that were consistent with empirical values (see **Figure 3d, S7 & Methods**). This analysis confirmed that the ‘diffuse’ PSD95 population does not correspond to complete spatial randomness. Instead, it aligns with a Thomas process fitting, indicating a degree of spatial order within the ‘diffuse’ population across all regions (see **Figure S8**). Taken together, we can conclude that most PSD95 is not localized within the PSD but still retains a small degree of spatial ordering on the nanoscale. We therefore decided to investigate the clustering of diffuse PSD95.

### PSD95 exists as multimers within the diffuse population

We have recently used single-molecule and super-resolution microscopy to analyze isolated PSD95 supercomplexes from brain homogenate, finding that the majority of PSD95 supercomplexes contain two or more copies of PSD95^25^. With our knowledge of the existence of the diffuse population here, we next sought to determine whether the individual PSD95 molecules were near each other, and therefore likely to reside as supercomplexes.

To achieve this, we revisited 2D PALM experiments, which gave higher lateral spatial resolution compared to the DHPSF^36^, and therefore allowed the direct observation of the spatial organization in the diffuse PSD95 subpopulation. The brain sections were immobilized on a glass coverslip and imaged *via* PALM using total internal reflection fluorescence (TIRF) microscopy (see **Methods**). Each PSD95-mEoS2 molecule was localized with a mean precision of 15.96 ± 2.91 nm (mean ± S.D., n = 5546 localizations). The acquired images had a mean resolution of 43.5 nm as determined by Fourier Ring Correlation^44^. A representative 2D SR image is shown in **Figure S9**. As with 3D data, clustering was used to distinguish the diffuse population of PSD95 from that residing in NCs. Using this approach, 70% of PSD95 was found in the diffuse population and using DBSCAN (radius of 125 nm, minimum of 10 localizations per cluster). Given that only half of the PSD95 proteins are fused to mEoS2 in heterozygous mice, and that not all mEoS2 will fold correctly, this suggests that the vast majority of PSD95 molecules in the diffuse population are close to a neighboring molecule and are therefore likely to be within the supercomplexes we have previously identified^25^.

Image averaging is a proven technique to increase the spatial resolution of SMLM images down to the level of 5 nm or higher^45^. We validated our image averaging analysis pipeline using TetraSpeck microspheres (see **Figure 4a**) and GATTAquant PAINT nanorulers (see **Figure 4b**), which together confirmed our ability to resolve objects separated by 40 nm. Next, we applied class averaging to 1402 supercomplexes containing two PSD95-mEoS2 proteins to investigate their spatial arrangement (see **Figure 4c**). This revealed that most PSD95 molecules had a peak-to-peak separation of ∼40 nm (see **Figure 4d**, 1402 dimers). However, a subset displayed a larger separation, resulting in a broad ‘tail-like’ distribution in the class averaged image. Together, **Figures 4d & 4e** show the distribution of spatial separations for both nanoruler and PSD95 images side by side. Further quantitative analysis revealed that 72% of PSD95 dimers were separated by a distance of 39 ± 2 nm, and 28% PSD95 dimers were separated by a larger distance of 94 ± 27 nm. The shorter separation distance is close to the spatial resolution that was achievable using PALM, and so we cannot rule out shorter separation distances that we have previously measured using MINFLUX in brain homogenate^25^. These data are not consistent with higher order geometries, such as trimers and tetramers, from the absence of their characteristic peak patterns of spatial separations (see **Figure S10**).

**Figure 4.**
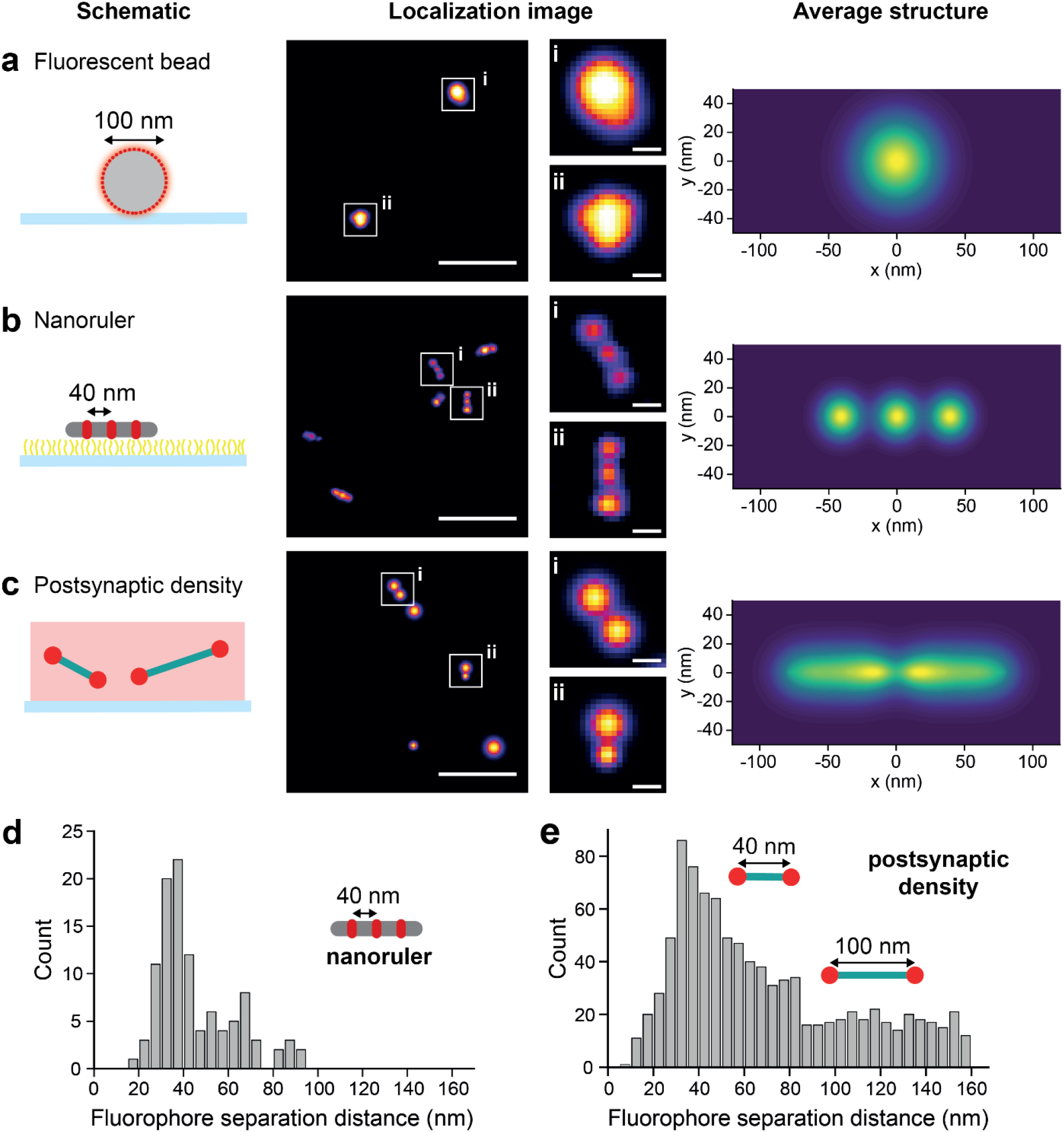
Identification of PSD95 dimers in the diffuse population. **a** SR analysis of TetraSpeck fluorescent beads. An example super-resolution image and the class average of 2534 beads is shown. **b** Super-resolution imaging of DNA-PAINT-based GATTAquant nanorulers. The nanorulers have three fluorophores, each spaced 40 nm apart. The class average of 828 trimeric nanoruler objects is shown. **c** PALM-based imaging of PSD95 endogenously tagged with mEos2 in tissue sections from the CA1 region of the hippocampus. The class average of 1402 dimeric objects is shown. These dimeric objects represent 15% of the total number of clusters present. Histogram of separation distances between fluorophore positions on **d** the GATTAquant DNA nanorulers and **e** the dimeric PSD95 clusters seen in tissue sections. Scale bars are 400 nm (large) and 40 nm (insert).

Given the existence of dimers, we next validated their spatial dimensions using a complementary labeling strategy. This strategy incorporated a HaloTag conjugated to a bright organic fluorophore to afford higher spatial localization precision for dual-labeled dimeric supercomplexes compared to mEos2. However, a limitation of this labeling strategy was the degree of dual-labelled dimers compared to that achievable with endogenous labeling. Photoactivatable Janelia Fluor 549 was selected for these experiments, which also revealed the existence of two spatially separated PSD95 populations at ∼40 nm and ∼100 nm, in agreement with our mEos2-PALM experiments (see **Figure S11**).

## Discussion

We demonstrate, to our knowledge, the first implementation of DHPSF SR microscopy in tissue sections, showing that the technique can super-resolve single fluorescent proteins in 3D, deep within brain tissue. Use of the DHPSF enabled us to enhance our understanding of the spatial organization of PSD95 in brain tissue. This approach confirms the presence of NCs of PSD95 in the PSD of excitatory synapses previously observed using 2D SMLM. Although the PSDs are the most obvious structures in the tissue sections, an abundance of PSD95 is found in diffuse clouds outside of these, most likely within the dendritic tree. This population is observed in both the CA1 and DG, which are populated by pyramidal neurons and granule cell neurons, respectively, suggesting that the diffuse population is found in the major classes of excitatory neurons and is not neuron-type-specific.

To ensure that the diffuse population was not an artifact of the mEos2-labeled PSD95, we imaged genetically modified mice expressing PSD95 fused to either eGFP or HaloTag using spinning disk confocal microscopy. Both imaging approaches confirmed the presence of PSD95 outside the PSDs; however, the proportion of diffuse PSD95 was lower than that observed with mEos2 labeling. This difference could be attributed to the lower detection efficiency of spinning disk confocal microscopy, or the poorer fitting of double helices in high-density/high-background PSD regions, which may exclude some PSD95 molecules. The finding that the majority of protein resides outside of the PSD can be rationalized by considering that the total volume of PSDs is minuscule: a simple back-of-the-envelope calculation estimates that the total volume of all PSDs is ∼0.02% of the entire mouse brain (see **Section S2**). While we cannot entirely rule out the possibility that a minority of these localizations may have been misidentified and belong to NCs within PSDs, their spatial uniformity throughout the imaging volume strongly suggests that the majority are not associated with PSDs. Additionally, there is no observed increase in density of these signals around the PSDs, further supporting this conclusion.

Further examination of the diffuse population identified a lesser degree of clustering, which we subsequently assigned as PSD95-containing supercomplexes, the majority of which contain two units. This finding supports biochemical studies on those purified from the mouse brain homogenate^13,26^. We have also recently used single-molecule and super-resolution microscopy to demonstrate that the majority of supercomplexes isolated in mouse brain homogenate contain two units of PSD95^25^. Furthermore, we showed that in addition to the entire supercomplex being turned over, protein can be degraded and replaced within each supercomplex, a finding that could have ramifications for long-term memory storage by proteins that have a lifetime ranging from hours to months^16^. In this study, we identified the presence of supercomplexes outside the PSD, although whether protein exchange occurs in this location remains unclear.

We also showed that the diffusely distributed supercomplexes containing two units form two populations with a spatial separation of 39 ± 2 nm or 94 ± 27 nm, which may be indicative of a differential composition of interacting proteins. Although we could not resolve supercomplexes within the densely packed NCs in the PSD, biochemical studies indicate that they also comprise two PSD95 units, leading us to speculate that PSD95 forms complexes in dendrites that are transported into synapses where they pack together into the NCs of the PSD.

Our results indicate that while the dense packing of complexes within the PSD enables PSD95 to be visible by confocal microscopy, the single molecule sensitivity of the DHPSF enabled the much larger diffuse population to be resolved and quantified and not manifested as low level autofluorescence. Therefore, the application of DHPSF and other multidimensional SR microscopy methods could further our understanding of whether other synaptic proteins are more widely distributed than previously known.

Mice carrying genetically tagged PSD95 have enabled the most comprehensive analysis of the spatial organization and distribution of any synaptic protein to date, spanning scales from single molecules to the whole brain^16,25,46–48^. These studies have revealed how PSD95 undergoes a hierarchical assembly of increasing molecular complexity from proteins, complexes, supercomplexes, and nanoclusters to form the mature and functional postsynaptic terminal of excitatory synapses. Moreover, the differential distribution of these supramolecular structures into different synapses on the dendritic tree, and between neuron types, produces a remarkable spatial diversity of synapses described by the synaptome architecture of the brain^15,17^. The stability of the synaptome architecture of PSD95-expressing synapses in the adult brain, despite PSD95 turnover, suggests that the regulation of the synaptic and extrasynaptic protein pools, as described in this study, is tightly regulated^16,17^. Our approaches, which are applicable to any synaptic protein, lay the groundwork for exploring the dynamics of PSD95 supercomplex populations, their mechanisms of transport, and assembly into synapses.

## Data and Code Availability

Supplementary analysis code and raw datasets are available at: 10.5281/zenodo.13332752^49^.

## Statistics and Reproducibility

No statistical methods were used to predetermine sample size. Where relevant, sample sizes are reported in the figure legends, and all experiments were performed independently at least three times to ensure results were repeatable.

## Acknowledgements

K.M. and M.H.H. acknowledge funding from UCB Biopharma and Dr Jim Love. The work was also funded by a Royal Society University Research Fellowship (UF120277 and URF\R\180029, to S.F.L.) and the EPSRC (RF1943723, to A.H.).

## Contributions

S.F.L, M.H, and S.G. conceived the project. S.D, K.M., E.B, A.H, C.A, and T.K performed the experiments. S.D, K.M, A.H, and C.A performed analysis of the localization data. E.B and S.G. provided the tissue sections and care and oversight for the mice used in the study. L.M and K.O.H wrote the PCF analysis code. A.K and A.S wrote the vLUME analysis code. G.C wrote the quantitative localization analysis code. C.A. and T.K. analyzed the confocal imaging data. S.G, M.H and S.F.L supervised the research. S.D, E.B, M.H and S.F.L wrote the manuscript with input from all the authors.

