## Supplementary Information for "3D super-resolution imaging of PSD95 reveals an abundance of diffuse protein supercomplexes in the mouse brain"

[1]S.D. and E.B. contributed equally to this work.
[2]To whom correspondence may be addressed.

### Table of contents

| | | |
|---|---|---|
| S1. | Materials and methods | 3 |
| S2. | Estimation of PSD volume to total brain volume | 8 |
| S3. | Supplementary figures | 9 |
| S4. | Description of supporting movies | 15 |
| | References | 16 |

#### S1: Materials and methods

**Tissue sectioning**. Animal procedures were performed in accordance with UK Home Office regulations and approved by Edinburgh University Director of Biological Services. Generation and characterization of PSD95-mEos2 fluorescent knock-in mouse line was described previously[1]. Adult 2- to 3-month-old heterozygous PSD95-mEos2 (n = 3) and wild-type C57BL/6 (n = 1) mice were fully anaesthetized by intraperitoneally injecting pentobarbital (0.10 mL, Euthatal) and, following the opening of the thorax, were perfused with PBS (1 × 12 mL) and post-fixed with 4% paraformaldehyde (PFA; #043368.9M, Thermo Scientific). The brains were dissected and placed in 4% PFA and incubated at 4°C for 3–4 hours. 4% PFA solution was replaced with 30% sucrose solution and the brains were left at 4°C for 48-72 hours. Brain tissue was embedded in OCT embedding matrix (CellPath) inside a plastic mould (#1003684132, Sigma-Aldrich) and frozen with liquid nitrogen and isopentane. Brains were stored at -80°C for up to 1 month. Glass coverslips (#1; 631-0142, VWR) were plasma cleaned and coated with poly-D-lysine (PDL, 1 mL, 0.22 µm filtered) for 30 min. Coverslips were then washed with distilled water ($dH_2O$). Fluorescent nanodiamonds (0.1 µm, 50 µL, nitrogen vacancy >900 NV/particle, Sigma-Aldrich) were applied and incubated for 30 min. The coverslips were then washed with $dH_2O$. Frozen brain samples were cut at 6 µm thickness using a cryostat (NX70 Thermo Fisher) to obtain coronal brain sections referring bregma level between -1.94 and -2.46 mm, capturing dorsal hippocampus. Cut brain sections were placed on the coverslips. A drop of $dH_2O$ was placed on a glass slide prior to picking up the brain sections to ensure the brain tissue laid flat. After cutting, brain sections were left to dry in the dark at room temperature overnight and were then stored at -80°C for up to 1 month.

**Double helix point spread function (DHPSF) microscopy.** The DHPSF is a custom widefield microscopy platform that incorporates additional optical elements to transform the shape of the standard PSF to encode the axial position of single emitters. This was built incorporating a 1.27 NA water-immersion objective lens (Plan Apo VC 60×, Nikon, Tokyo, Japan) to image above the coverslip surface. The DHPSF transformation was achieved by introducing additional optics into the emission path of a conventional fluorescence microscope (Eclipse Ti-U, Nikon) with the objective lens mounted onto a scanning piezo stage (P726 PIFOC, PI, Karlsruhe, Germany). A 4*f* system of lenses placed at the conjugate image plane relayed the image onto an EMCCD detector (Evolve 512 Delta, Photometrics, Tucson, AZ, USA; 16 µm pixel size). A double-helix phasemask (PM) (DoubleHelix, Boulder, CO, USA; 580 nm optimized) placed in the Fourier plane of the 4*f* system performed the DHPSF transformation. Excitation and activation illumination was provided by 561 nm (200 mW, CoboltJive 100, Cobolt, Solna, Sweden; ~0.25 kW $cm^{-2}$) and 405 nm (120 mW, iBeam smart-405-s, Toptica, Munich, Germany; <7 W $cm^{-2}$) lasers, respectively. The lasers were circularly polarized, collimated, and focused to the back aperture of the objective lens. The fluorescence signal was then separated from the excitation beams into the emission path by a quad-band dichroic mirror (Di01-

R405/488/561/635-25×36, Semrock, Rochester, NY, USA) before being focused at the image plane by a tube lens (f = 200 mm, Nikon). Finally, long-pass and band-pass filters (BLP02-561R-25 and FF01-580/14-25, Semrock) placed immediately before the camera isolated the fluorescence emission. An exposure time of 50 ms was used to image the PSD95-mEos2.

The conversion gain of the EMCCD detector (Evolve 512 Delta, Photometrics) for DHPSF microscopy was measured frequently using a custom MATLAB script (https://github.com/TheLeeLab/cameraCalibrationCMOS). The in-built 'Rapid-Cal' function also served to calibrate the EM gain register on a weekly basis. The CCD gain value was measured at 0.154 counts/photoelectron which—when operated at an EM gain of 250—gave a total conversion gain value of 38.5 counts/photoelectron.

**Preparation of fluorescent bead samples for DHPSF calibration.** Glass slides (VWR, 631–1570) were cleaned under argon plasma (PDC-002, Harrick Plasma, Ithaca, NY) for 1 hour and incubated with poly-L-lysine (PLL, 50 µL, 0.1% w/v, Sigma-Aldrich, P820) for 30 minutes. Glass slides were then washed with PBS (3 × 50 µL) and incubated with fluorescent beads (50 µL, 4.6 × $10^7$ particles $mL^{-1}$; 0.10 µm, TetraSpeck microspheres, Thermo Fisher, Waltham, MA, USA) for 2 mins before washing further with PBS (3 × 50 µL). The sample was then transferred to the microscope for imaging. Here, the piezo-mounted objective was scanned axially through the sample in 60 nm steps over 4 µm, recording 10 frames of 30 ms exposure at each step. This calibration file was subsequently used for 3D reconstruction using easy-DHPSF fitting software[2].

**DHPSF localization analysis.** Initially, ImageJ was used to remove autofluorescence background from raw localization datasets by subtracting the average intensity offset of the z-stack. The immobile fiducial marker was therefore processed separately since it was steadily fluorescent. Super-localization of the DHPSF was conducted using easy-DHPSF, which has been described previously[2]. In brief, single-molecule fluorescence was identified using template matching and localized in 3D by double Gaussian fitting of the DHPSF using non-linear least squares minimisation. The lateral position of each fluorophore was determined from the midpoint of both Gaussian lobes, while the z position was calculated from the angle between the two Gaussian lobes using the calibration curve produced from fluorescent beads. The fitting precision of the microscope platform has been evaluated previously at Ref. [3] (Figure 1b therein) using the following equation:

$$\text{Total 3D precision} = \sqrt{\Delta x^2 + \Delta y^2 + \Delta z^2}$$

where $\Delta x$, $\Delta y$, and $\Delta z$ refer to the standard deviation in the position of a point emitter at a given photon flux. This gave a total 3D precision of 38 ± 8 nm for mEos2 (see **Figure S1**).

To avoid overcounting errors the inherent photophysical properties of mEos2 were evaluated for DHPSF microscopy. Prior to cluster analysis, the proportion of repeated blinks from the same fluorophore was determined experimentally. The results are presented in Figure S1, which show that the probability of detecting a recurring signal from the same fluorophore within a sphere of 50 nm radius is 2% over a 50-frame sliding window. This is lower than typically observed for mEos[4] but can be rationalised by to the low photon-efficiency of the DHPSF phase mask, which reduces overall signal by approximately 20–30%[5], and the higher laser powers used in this work. Therefore, the risk of overcounting molecules was considered negligible.

**Fiducial correction for DHPSF microscopy.** Drift correction was performed by localizing the position of a fiducial marker in each frame and subtracting the resulting 3D points from all localizations in the corresponding frame. Fluorescent nanodiamonds (10 µg $mL^{-1}$ in $dH_2O$; 100 nm, nitrogen vacancy >900 NV $particle^{-1}$, Sigma-Aldrich) were used as fiducial markers to account for 3D focal drift. Microscope slides were incubated with PLL (50 µL, 0.1% w/v, Sigma-Aldrich, P820) for 30 min and washed with PBS (3 × 50 µL) before adding the nanodiamond solution. The sectioned brain was then adhered to the coated slides and the nanodiamonds left to diffuse through the sectioned tissue to reside at different depths.

**Nanocluster analysis.** The size, volume and number of localisations per cluster were calculated and visualised using vLUME desktop software (see Figure 2e–g)[6]. First, the NC and diffuse populations were segmented using DBSCAN (radius = 125 nm, $N_{localisations} \geq 10$). Radius and volume were determined using bespoke analysis scripts in vLUME that implemented triple product convex mesh fitting to calculate the volume of a parallelepiped (composed of six equal tetrahedra). In brief, the cross product was used to find the area of the parallelogram formed by two vectors. The volume was then obtained by projecting the diagonal length onto the base's normal (which is implicitly done via the dot product) as the cross-product’s area vector served as the scaled normal. The resulting volume was either convex or concave, and was approximated by measuring a spherical volume with a radius equal to the distance from the centroid to the furthest vertex.

**Pair correlation function analysis.** First, the nanocluster (NC) and diffuse populations were segmented using a mixture model applied to the k-nearest neighbour (nn = 10) distance computed for each point (as implemented by the *nnclean* function in the R *spatstat* library).

Pair Correlation Function (PCF) analysis was then implemented to determine the extent of clustering for the NC, PSD and NC/PSD distributions using three fitting algorithms[7]. The first algorithm comprised a complete spatial randomness test (CSR, see **Figure S6a**) for which simulated datasets of random points were generated with equal volume and localisation density as the raw data (N = 49). The second was a single Thomas fit (Neyman-Scott cluster process[8], where the clusters follow 3D Gaussian distributions around parent points, see **Figure S6b**). This process generated simulated datasets of clustered localisations where parameters were informed by experimental data (125 nm radius, 10–50 localisations per cluster, N = 49). Finally, a double Thomas fitting algorithm produced simulated cluster-of-clusters datasets consistent with the experimentally observed NC population only (see **Figure S2c**). For each population, a model selection and a parameter estimation step were applied.

Both model selection and parameter estimation were based on the PCF used, a summary statistic with a known analytical expression for the three models above. A Poisson process (describing CSR) has a $g(r) = 1$. A modified Thomas process has a $g_1(r)$ described by Equation S1

$$g_1(r) = 1 + \frac{1}{\kappa}(4\pi\sigma_1^2)^{-3/2}\, exp\left(\frac{-r^2}{4\sigma_1^2}\right) \tag{S1}$$

and a double Thomas process has a $g_2(r)$ described by Equation S2

$$g_2(r) = 1 + \frac{1}{\kappa\mu}(4\pi\sigma_1^2)^{-3/2} exp\left(\frac{-r^2}{4\sigma_1^2}\right) + \frac{1}{\kappa}(4\pi(\sigma_1^2 + \sigma_2^2))^{-3/2} exp\left(\frac{-r^2}{4(\sigma_1^2+\sigma_2^2)}\right) \tag{S2}$$

where $\kappa$ is parent intensity.

Fitting the experimental PCF to the theoretical description allowed for estimation of the model parameters. For the Thomas process this included cluster size (σ) and cardinality (μ). For double Thomas process this includes the cluster sizes ($\sigma_1$ and $\sigma_2$) and cardinalities (μ and ν). In order to select the best model, envelope tests were performed (N = 49 replicates) and the envelope of the PCF summary statistics were computed. The empirical PCF was superimposed on the envelope graphs and the overlap indicates how well the model describes the experimental data.

In summary, the NC populations were fitted to Thomas and double Thomas processes, while the diffuse populations were fitted to CSR and Thomas processes. Representative code and datasets are available via the Zenodo repository associated with this work[7].

**2D PALM imaging.** Two-dimensional PALM experiments were performed on a home-built TIRF microscope described previously[9]. Briefly, collimated laser light at 405 nm (Cobolt MLD 405–250 Diode Laser System, Cobalt, Sweden) and 561 nm (Cobolt DPL561-100 DPSS Laser System, Cobalt, Sweden) was aligned and directed parallel to the optical axis at the edge of a 1.49 NA TIRF Objective (CFI Apochromat TIRF 60×C Oil), mounted on an inverted microscope (Ti2, Nikon). The microscope was fitted with a perfect focus system to auto-correct the z-stage drift. Fluorescence collected by the same objective was separated from the returning TIR beam by a dichroic mirror (Di01-R405/488/561/635, Semrock, Rochester, NY, USA), and was passed through appropriate emission filters (LP02-568RS, FF01-587/35, Semrock, Rochester, NY, USA). Fluorescence was then passed through a 2.5× beam expander and recorded on an EMCCD (Evolve 512 Delta, Photometrics, Tucson, AZ, USA; 16 μm pixel size) operating in frame transfer mode. This produced a pixel size at the sample plane of 103 nm. The CCD gain was measured at 0.0870 counts/photoelectron which, when operated with an EM gain of 250, gave a total conversion gain of 21.0 counts/photoelectron. Images were recorded with an exposure time of 50 ms. The microscope was automated using the open-source microscopy platform Micromanager[10]. Borosilicate glass coverslips (20 × 20 mm, VWR International) were cleaned under argon plasma (Zepto, Diener) for 30 min to remove any fluorescent residues. The tissue sections, prepared as above, were then added to the coverslips. The samples were illuminated with 15 cycles at 405 nm (75 W $cm^{-2}$, 1 s) and 561 nm (1.5 kW $cm^{-2}$, 10 s) irradiation until all molecules were photobleached. For the identification of PSD95 dimers in the diffuse population, GATTA-PAINT 40 nm nanorulers (PAINT 40R, fluorophore: ATTO 655) and TetraSpeck microspheres (100 nm, T7279) were used.

The data was analysed using the Peak Fit (GDSC SMLM, ImageJ plug-in) to output localizations of the individual fluorophores. A signal strength threshold of 20 and precision threshold of 40 nm were used. Focal drift was corrected for using the in-build Drift Calculator function in the GDSC SMLM plugin. A custom written MATLAB clustering script was used to classify localizations into clusters. All localizations were first ordered by precision from low to high. To correct for multiple localizations of the same fluorophore, the script combined localizations within the precision of another localization into one object. The distances between all objects were then calculated. Objects separated by less than 160 nm were grouped into clusters. The number of objects in each cluster were then counted. Clusters containing one object were defined as “monomeric”, clusters containing two objects were defined as “dimeric” and so forth. Information pertaining to the clusters (number of objects, x-y position of objects, average precision) was output as a text file.

**Class averaging of 2D data.** To produce the class averaged images, the centre point of each feature (PSD95 dimers, DNA nanorulers, or fluorescent beads) was found and the object rotated into the y = 0 plane. The average of all objects was then calculated and plotted as a surface plot, with the z axis representing the density of features in the x-y plane. The average precision of the localisations used in the class average was 25 ± 5 nm, which was determined from the photon budget of mEos2, and subsequently photon-matched for the beads and nanorulers.

**Preparation of histological sections.** Mice were anesthetized by intraperitoneal injection of dolethal (pentobarbital). The animal was then transcardially perfused with PBS (10 ml) followed by paraformaldehyde (PFA, 10 ml, 4% in PBS). The brain was dissected, kept in 4% PFA for 3 hours, and then transferred to 30% sucrose in PBS. After 48–72 hours of incubation at 4 $^{o}$C, the sample was embedded in OCT and frozen using liquid nitrogen. The samples were kept at –80 $^{o}$C until use. Frozen brains were cut at 18 mm thickness using a cryostat (CM3050 S, Leica) to obtain sagittal sections. The brain sections were placed on Superfrost Plus glass slides (Epredia) and left to dry up in the dark at room temperature overnight and kept at –20°C until use. As for PSD95-eGFP sections, the samples were incubated in PBS for 15 min and then embedded in MOWIOL solution with coverslips.

**Labeling brain sections with HaloTag ligand.** PSD95-Halo brain sections were first incubated in PBS for 10 min and then in PBS with/without JF552 HaloTag ligand (800 nM). The sections were washed three times with PBS containing 0.2% Triton X-100 and then twice with PBS without detergent. The labelled sections were embedded in MOWIOL solution with coverslips.

**Confocal microscopy.** Imaging was performed using spinning disk microscope (Nikon, Eclipse Ti2) equipped with a 100× objective lens. Images of 862 x 826 pixels in size and 16-bit depth were obtained. eGFP and JF552 were excited at 488 nm and 561 nm laser, respectively.

#### S2: Estimation of PSD volume to total brain volume

The average volume of the mouse brain is 415 $mm^3$, and the volume of each brain region is given in **Table S1**[11]. Data describing the density of excitatory synapses in each region was taken from Santuy *et al.* and is shown in the second column of Table S1[12]. The number of excitatory synapses in each region was calculated, along with the total number of excitatory synapses.

The volume of the PSD was calculated assuming the PSD is an oblate spheroid, with the area of the circular 2D projection being 0.12 $\mu m^2$ [13] and the thickness being 23 nm[14]. Using Equation S3 where $A$ is the area of the circular 2D projection and $r$ is the semi-minor axis of the oblate spheroid, the volume of the PSD was calculated to be $1.4 \times 10^{-21}$ $m^3$.

$$V = \frac{1}{3}\pi r \times A \tag{S3}$$

The total number of excitatory synapses in the mouse brain was calculated to be $4.4 \times 10^{11}$.

The total volume of PSDs in excitatory synapses was calculated to be $6.2 \times 10^{-11}$ $m^3$.

The volume of the mouse brain is $4.15 \times 10^{-7}$ $m^3$.

The volume of PSDs is therefore only 0.02% of the total brain volume.

**Table S1.** Information used in the estimation of PSD volume to total brain volume.

| **Region** | **Volume ($x10^{10}$ $\mu m^3$)**[11] | **Number of synapses per unit volume ($\mu m^{-3}$)**[12] | **Total synapses ($\times 10^{10}$)** | **Total volume of PSDs ($\times 10^7$ $\mu m^3$)** |
|---|---|---|---|---|
| Isocortex | 11 | 1.7 | 19 | 27 |
| Hippocampus | 2.3 | 1.8 | 4.1 | 5.7 |
| Striatum | 1.5 | 1.3 | 2.0 | 2.8 |
| Diencephalon | 4.0 | 0.79 | 3.2 | 4.5 |
| Brainstem | 5.0 | 0.26 | 1.3 | 1.8 |
| Cerebellum | 5.2 | 0.47 | 2.4 | 3.4 |
| Fiber tracts | 1.4 | 0.020 | 0.028 | 0.039 |
| Other | 11 | 1.1 | 12 | 17 |
| **Total** | **41** | **-** | **44** | **62** |

#### S3: Supplementary Figures

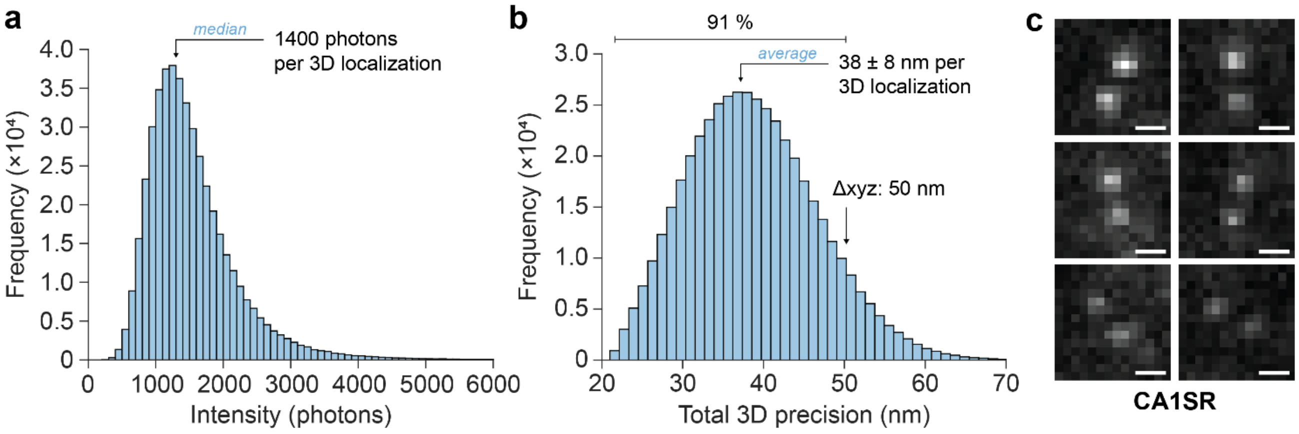

**Figure S1. Distribution of single-molecule intensities. a** Distribution of PSD95-mEos2 single-molecule intensities via DHPSF SR microscopy. A median of 1400 photons were detected per molecule in 3D in CA1SR region (representative of all regions) using easy-DHPSF software. **b** Empirically determined localisation precision using data from Ref 3 (Figure 1b, same microscope). An average 3D precision of 38 ± 8 nm was determined. 91% of localizations were below 50 nm precision. **c** Raw DHPSF single molecule localisation data illustrating the range of signal-to-noise ratio obtained.

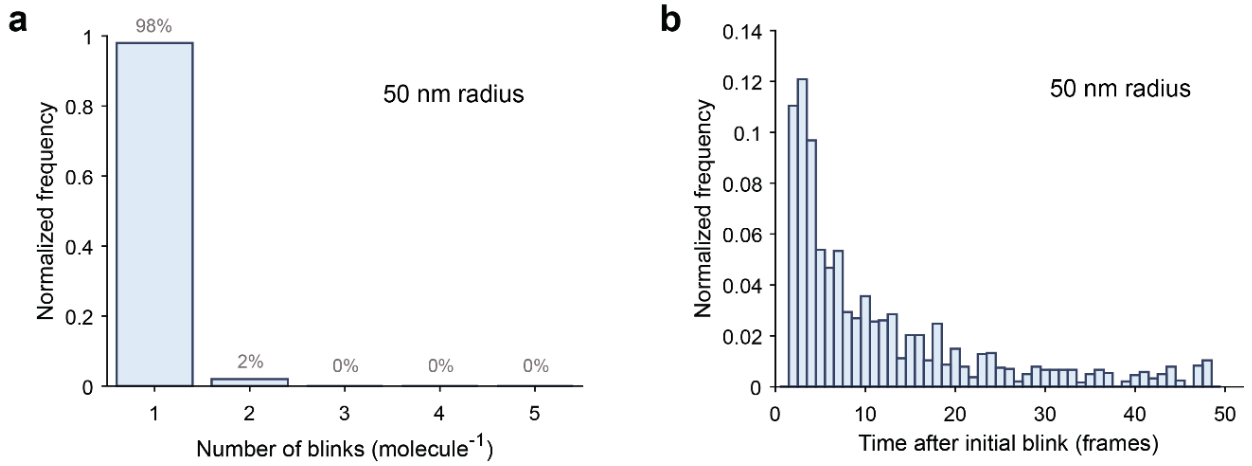

**Figure S2. Effect of repeat photo-blinking from a single FP on analysis. a** The optimisation of frame rate, illumination power density and photon efficiency of the DHPSF with mEos2 meant that 98% of all fluorescent events were captured within a single frame. **b** The 2% of molecules that blinked twice were analysed temporally within a 50-frame window. For this subset of mEos2 molecules, a second blink was detected from 51% of molecules within 8 frames following the initial blink. A radius of 50 nm was selected based on the precision histogram in **Figure S1b**.

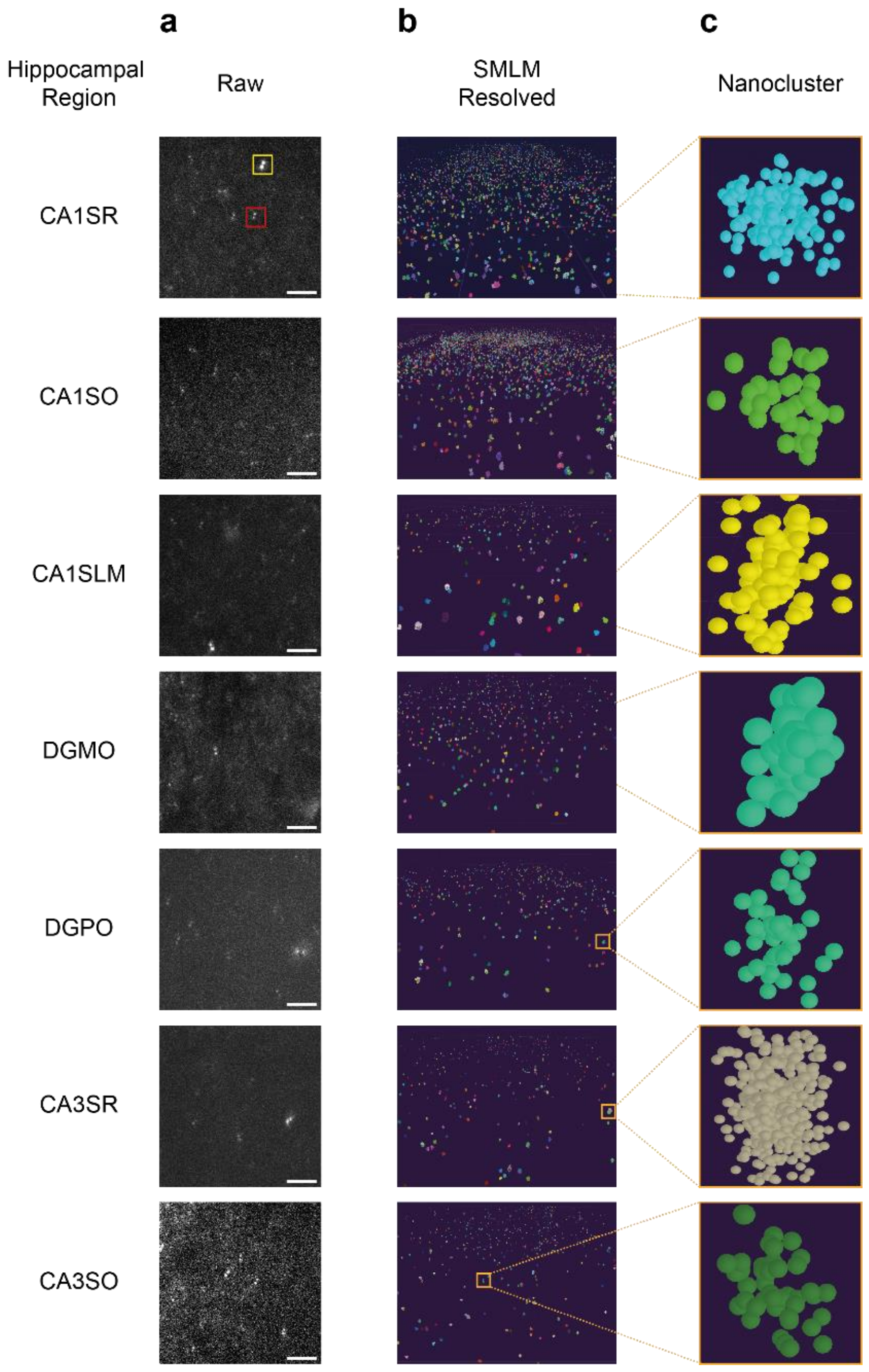

**Figure S3. Example DHPSF data collected from different brain regions.** Seven examples of collected data, with super-localised reconstructions and an example of a single nanocluster. **a** The seven hippocampal regions studied in this paper (CA1SR, CA1SO, CA1SML, DGMO, DGPO, CA3SR, CA3SO) and a representative frame of raw DHPSF data from each of these regions. Highlighted in the first box (CA1SR) are the fiducial marker (yellow) and a single PSD95-mEos2 localisation (red). Scale bar is 10 µm. **b** The corresponding super-resolved data visualised in vLUME from 50,000 frames in each hippocampal region. Each dataset shows the PSD95 nanoclusters imaged in the hippocampus represented as demarcated colours. Each region is approximately 100 × 100 × 4 µm. **c** Example of a single PSD95 nanocluster (maximum radius of 125 nm) in each hippocampal region. CA1SR, CA1 stratum radiatum; CA1SO, CA1 stratum oriens; CA1SLM, CA1 stratum lacunosum-moleculare; DGMO, dentate gyrus molecular layer; DGPO, dentate gyrus polymorph layer; CA3SR, CA3 stratum radiatum; CA3SO, CA3 stratum oriens.

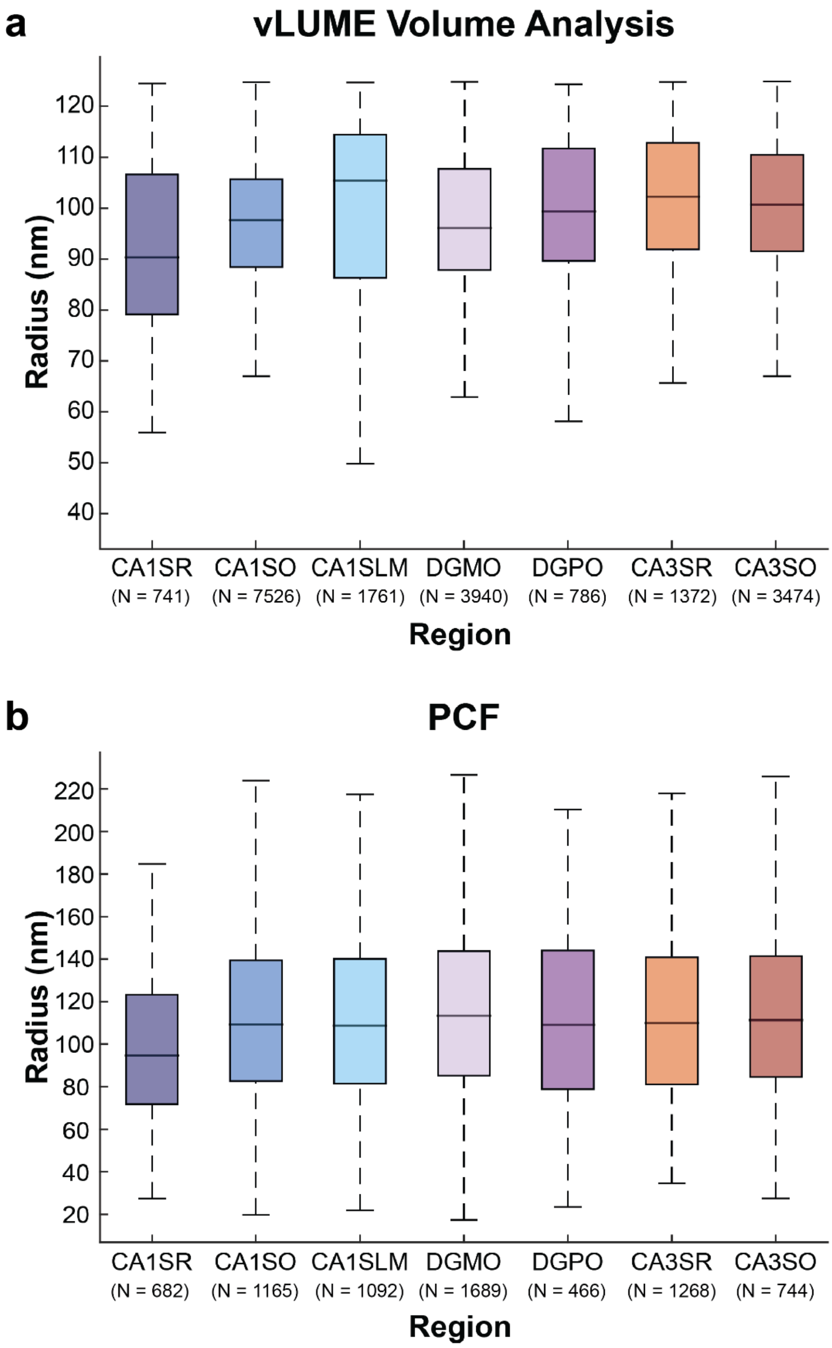

**Figure S4. Radii of nanoclusters determined using different approaches. a** The nanocluster radius size by region. This radius size was calculated from the volume box plots in Figure 2e. The volume has been approximated to be spherical to calculate its radius. **b** This is complemented by nanocluster radius sizes calculated by Pair Correlation Fit. Box plots represent the median value and interquartile range, where whiskers denote the lower and upper limits not considered outliers. Number of nanoclusters is denoted by N for each box plot.

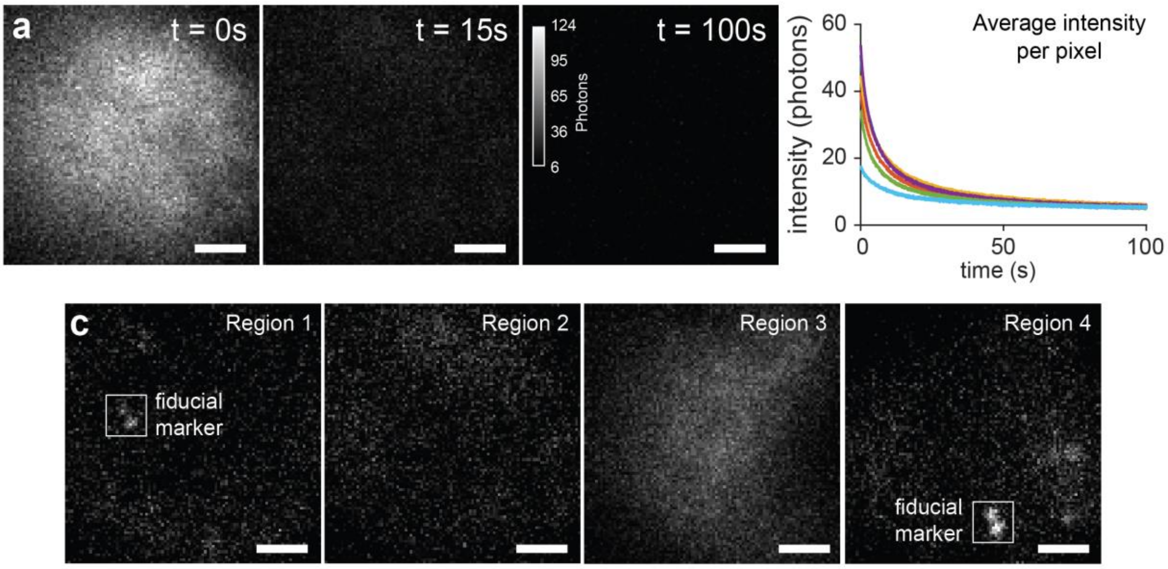

**Figure S5. Negative control of WT brain tissue. a** Representative images of wild-type (WT, no mEos2-PSD95) brain tissue used as a negative control under identical imaging conditions to experimental samples (see **Methods**). This control confirms that background fluorescence was almost completely bleached after 15 s (300 frames, 50 ms exposure) of illumination. During data acquisition in this work, the sample was first bleached for this duration. Scale bar is 5 µm. **b** A plot of average pixel intensity over a 100 s (2000 frame) period. Colours represent repeats in different regions of the hippocampus (N = 6). **c** No false positive localisations were detected in the WT brain tissue as shown by the representative frames. Fluorescent nanodiamonds (FNDs) used a fiducial markers are indicated with white boxes, which are expanded on the right as a z-projection over 2000 frames (100 s) to confirm their constant emission.

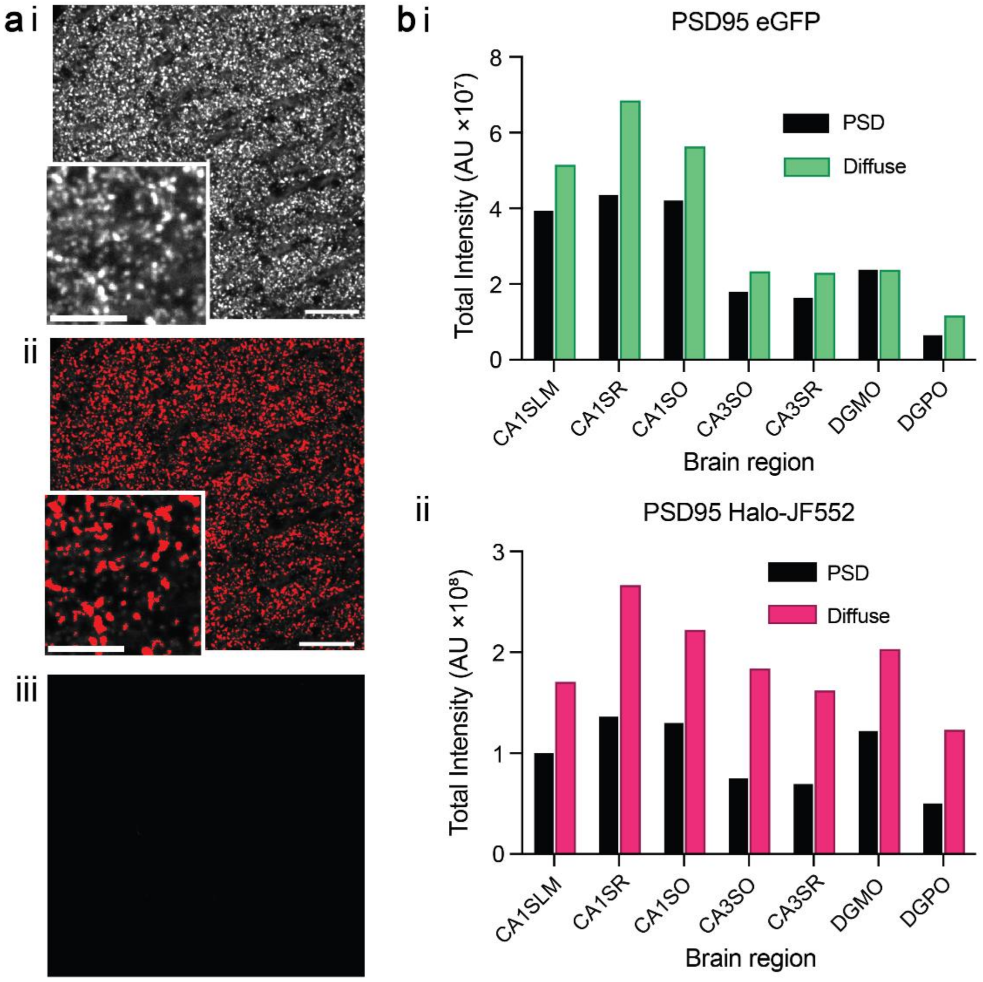

**Figure S6. Quantification of PSD95 found in the PSD or as diffuse protein using spinning disk confocal microscopy. ai** Representative image of hippocampal sub region (CA1SR) of a PSD95-Halo mouse (Ref. [15]) labeled with a JaneliaFluor-552 (JF552) HaloTag ligand and imaged using spinning disk confocal microscopy. Scale bar 100 µm, inset scale bar 5µm. **aii** Same field of view showing segmentation of PSDs. PSDs are defined by applying a threshold using Otsu's method. Intensities above the threshold were classified as PSDs, while diffuse regions were classified as areas below the threshold. **aiii** Hippocampal sub-region (CA1SR) of an unlabeled PSD95-Halo mouse (contrast-matched to **ai**). **b** Total intensities from PSDs and diffuse regions across different hippocampal subregions taken from an **i** PSD95-eGFP mouse (Ref. [13]), and **ii** PSD95-Halo mouse labeled with a JF552 HaloTag ligand. The total intensities for PSD (Black) and diffuse (Green eGFP/Magenta JF552) regions were background-subtracted based on the average of corresponding unlabelled brain regions. This analysis highlights the dominant contribution to the total intensity from a diffuse population of PSD95 outside of the PSD as well as the variation of populations across hippocampal regions. Homozygote of PSD95-Halo and heterozygote of PSD95-eGFP were used for the experiment.

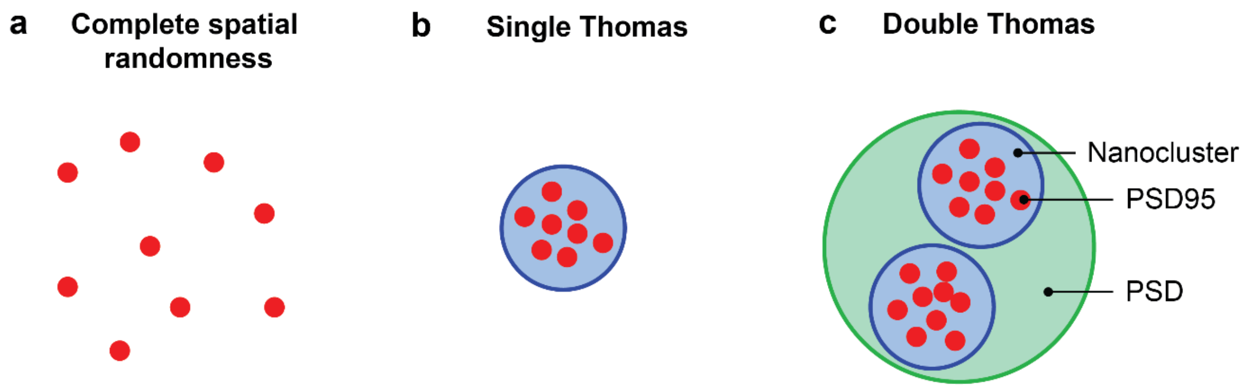

**Figure S7. Pair correlation fit analysis schematic.** Three different methods of clustering analysis are presented. **a** Localisations are completely spatially random. **b** Single Thomas process outputs nanocluster size and the extent of clustering. **c** Double Thomas process outputs the size of a nanocluster, a PSD, and the extent of clustering for each.

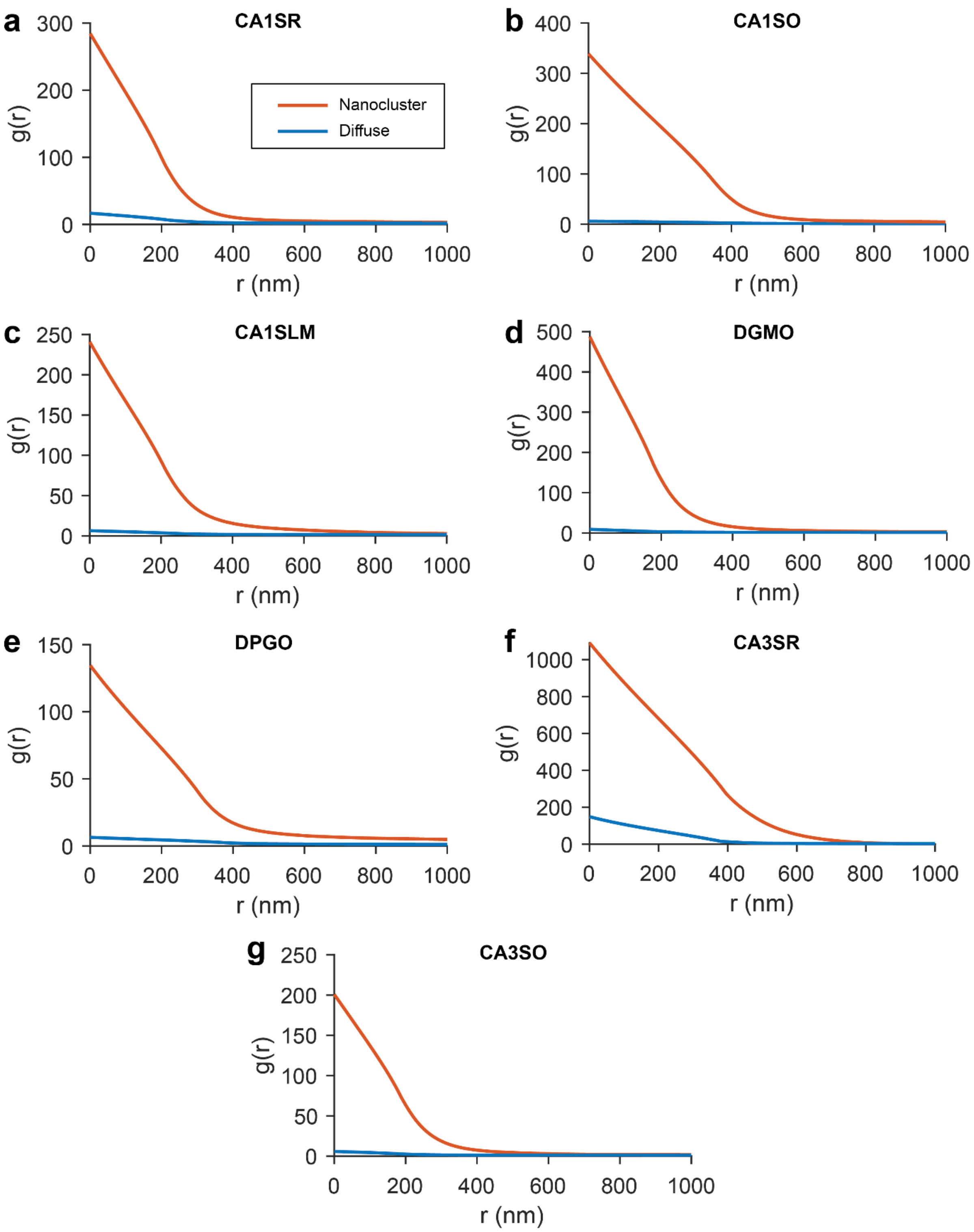

**Figure S8. Pair correlation of diffuse and nanocluster populations in different brain regions.** PCF fits for each hippocampal sub-region showing diffuse (blue) and NC (orange) fittings using single Thomas fitting. **a** CA1SR **b** CA1SO **c** CA1SLM **d** DGMO **e** DGPO **f** CA3SR **g** CA3SO.

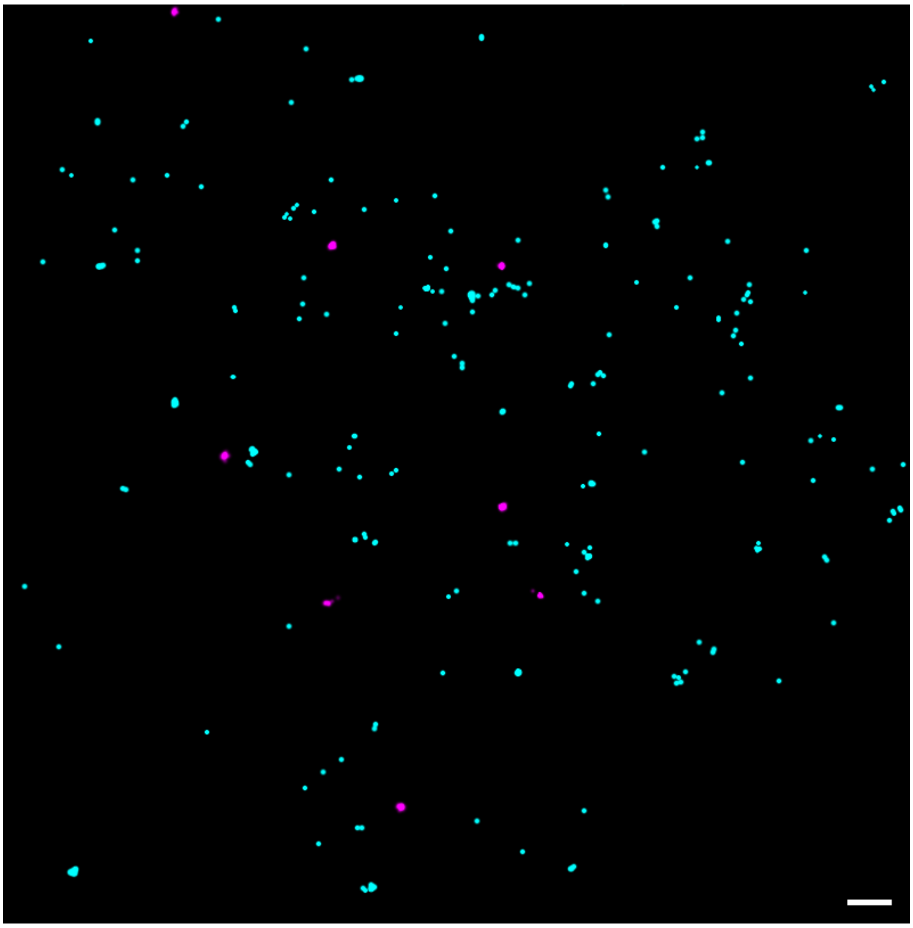

**Figure S9. 2D PALM images of PSD95-mEoS2 localizations in murine brain tissue**. Clustered (magenta) and non-clustered localizations (cyan). Data were generated using the Peak Fit (GDSC SMLM, ImageJ plug-in) to output localizations of the individual fluorophores. A signal strength threshold of 20 and precision threshold of 40 nm were used. A custom written Python script was used to implement a DBSCAN algorithm classify localizations into clusters, with parameters ε and minimum localizations set to 1.0 and 10 respectively. A total of 2753 localizations were clustered and 5546 non-clustered, indicating an extra-synaptic population of around 70%. Scale bar = 500 nm.

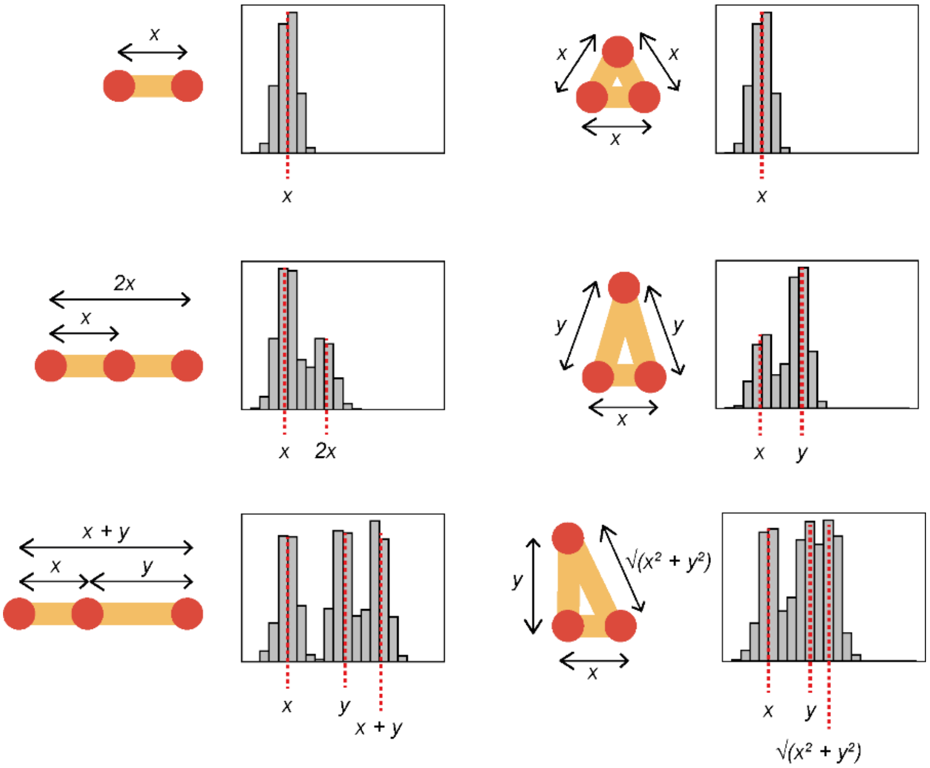

**Figure S10.** Schematic showing possible geometries of PSD95 clusters that would produce two distinct distance populations. None of these or higher order geometries represent the data we collected. Therefore, there must be two populations of dimeric clusters with distinct separation distances.

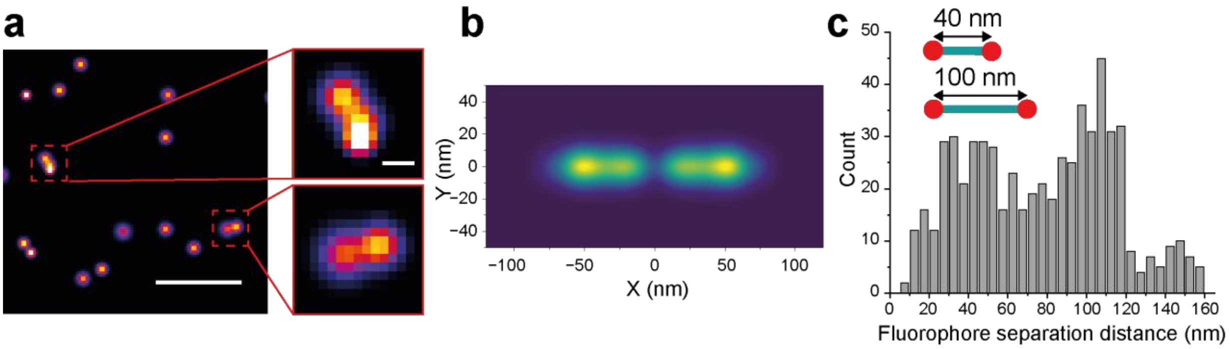

**Figure S11. Identification of dimers in PSD95 endogenously fused to HaloTag and labelled with photoactivatable Janelia Fluor 549 (PAJF549). a** Example PALM image of PSD95 endogenously tagged with HaloTag conjugated to PAJF549 in tissue sections from the CA1 region of the hippocampus. Scale bar = 400 nm, Inset scale bar = 40 nm. **b** The class average of 602 dimeric objects is shown. **c** Histogram of separation distances between the dimeric PSD95 clusters seen in tissue sections.

#### S4: Supporting Movies

**SI Movie 1 | PSD95-mEos2: Raw DHPSF localization data.**

Representative raw single-molecule localisation data. An exposure time of 50 ms was used with power densities of 0.25 kW $cm^{-2}$ and 0.7 W $cm^{-2}$ for the 561 and 405 nm laser lines, respectively.

**SI Movie 2 | PSD95-mEos2: Diffraction limited vs. super-resolved.**

Representative diffraction-limited image vs. super-resolution image of PSD95-mEoS2 taken from the CA1SR region.

**SI Movie 3 | PSD95-mEos2: Clusters and supercomplexes.**

Shows a representative super-resolution reconstruction of PSD95-mEoS3 taken from the CA1SR. The movie rotates around a central axis. Initially the reconstruction is shown with a density threshold applied to accentuate the PSD95 nanoclusters, this makes larger clusters more obvious and is similar to the analysis applied in our previous study. Halfway through the movie we switch to a rendering that shows all localisations (show in red). These data support the findings in Fig 3c that the vast majority of localisations do not reside in the NCs.
